# Immortalization and Functional Screening of Natively Paired Human T Cell Receptor Repertoires

**DOI:** 10.1101/2021.10.19.465057

**Authors:** Ahmed S. Fahad, Cheng Yu Chung, Sheila N. Lopez Acevedo, Nicoleen Boyle, Bharat Madan, Matias F. Gutiérrez-González, Rodrigo Matus-Nicodemos, Amy D. Laflin, Rukmini R. Ladi, John Zhou, Jacy Wolfe, Sian Llewellyn-Lacey, Daniel C. Douek, Henry H. Balfour, David A. Price, Brandon J. DeKosky

**Affiliations:** Department of Pharmaceutical Chemistry, The University of Kansas, Lawrence, KS 66044, USA; Vaccine Research Center, National Institute of Allergy and Infectious Diseases, National Institutes of Health, Bethesda, MD 20892, USA; Division of Infection and Immunity, Cardiff University School of Medicine, University Hospital of Wales, Cardiff CF14 4XN, UK; Department of Laboratory Medicine and Pathology, University of Minnesota Medical School, Minneapolis, MN 55455, USA; Department of Pediatrics, University of Minnesota Medical School, Minneapolis, MN USA; Systems Immunity Research Institute, Cardiff University School of Medicine, University Hospital of Wales, Cardiff CF14 4XN, UK; Department of Chemical Engineering, The University of Kansas, Lawrence, KS 66044, USA; Department of Chemical Engineering, Massachusetts Institute of Technology, Cambridge, MA 02142, USA; The Ragon Institute of MGH, MIT, and Harvard, Cambridge, MA 02139, USA

**Author notes:** These authors contributed equally (Ahmed S. Fahad, Cheng Yu Chung).

## Abstract

Functional analyses of the T cell receptor (TCR) landscape can reveal critical information about protection from disease and molecular responses to vaccines. However, it has proven difficult to combine advanced next-generation sequencing technologies with methods to decode the peptide-major histocompatibility complex (pMHC) specificity of individual TCRs. Here we developed a new high-throughput approach to enable repertoire-scale functional evaluations of natively paired TCRs. In particular, we leveraged the immortalized nature of physically linked TCRα:β amplicon libraries to analyze binding against multiple recombinant pMHCs on a repertoire scale. To exemplify the utility of this approach, we also performed affinity-based functional mapping in conjunction with quantitative next-generation sequencing to track antigen-specific TCRs. These data successfully validated a new immortalization and screening platform to facilitate detailed molecular analyses of human TCRs against diverse antigen targets associated with health, vaccination, or disease.

## Introduction

T cells form an essential component of adaptive immunity. Each human possesses billions of T cells, each of which expresses a somatically rearranged heterodimeric T cell receptor (TCR). In turn, each TCR recognizes a unique array of peptides bound to cell surface-expressed major histocompatibility complex (pMHC) molecules^1^. A detailed molecular understanding of CD4^+^ and CD8^+^ T cell responses is critically important for continued progress in human immunology and the rational development of immune-based therapies and vaccines. However, comprehensive TCR studies are complicated by the fact that each individual expresses up to six different MHC class I proteins (two allotypes each for HLA-A, HLA-B, and HLA-C) and up to six different MHC class II proteins (two allotypes each for HLA-DP, HLA-DQ, and HLA-DR), all of which bind distinct subsets of the universal peptidome. The TCR specificity landscape has been estimated to exceed a theoretical diversity of 10^15^ (ref^2^), and methods to decode pMHC antigen specificities of the patient-specific TCR repertoire will catalyze new scientific insights and disease treatments.

It is currently impossible to analyze all potential interactions that could occur in any given individual between pMHC molecules and TCRs. Most technologies to identify and monitor antigen-specific TCR sequences rely on screening limited numbers of viable T cells in blood or tissue samples. Moreover, the composition of T cell libraries can be altered by repeated *in vitro* expansion, which is necessary for studies of antigen specificity^3^, and methods based on limiting dilution or the isolation of single cells are generally restricted to small-scale analyses of TCRα:β pairs^4–7^. The difficulties associated with maintaining and studying primary human T cells *in vitro* have also prevented functional screening against large panels of different pMHC antigens. In addition, fate decisions and functional responses are largely dictated by TCR affinity in the context of any given pMHC, and accordingly, technologies that enable the quantification of TCR affinity across multiple pMHCs are highly desirable^8–10^.

Several new approaches to high-throughput TCR analysis have emerged in recent years. Impressive efforts have leveraged the power of single-cell transcriptomics to identify increasing numbers of antigen-specific clonotypes, but these studies are still limited by the labile nature of primary human T cells and the inability to perform repeated functional interrogations against TCR libraries^11–13^. Improved high-throughput technologies are therefore needed to characterize the scope of TCR interactions against pMHC antigens. Mass cytometry enables screening against large panels of metal-conjugated pMHCs, but fails to capture functional connections with individual TCR sequences^14–17^. Oligonucleotide tagging methods have been used similarly to identify multiple pMHC specificities^18^, but single-cell binding information is generally limited to relatively small numbers of TCRs^19^. Accordingly, single-cell droplet RNA sequencing has emerged as a rapidly growing platform for TCRα:β discovery^20–23^. Although a major advance, droplet-based methods are incompatible with TCRα:β gene immortalization, which precludes renewable functional screening and high-throughput affinity assays against pMHC panels.

High-throughput functional screening is critical for the identification of antigen cross-reactivity mediated by degenerate TCRs. Cross-reactivity has been associated with enhanced immune protection against variable pathogens^24–26^, but cross-reactivity can also lead to adverse patient outcomes in the context of immunotherapeutic TCRs^27^. Natively paired TCRα:β chains have been recovered from single cells and expressed in reporter cell lines to analyze functional reactivity^7,20,28,29^. However, these approaches are limited in scope, and library-scale methods that enable affinity-based screening are urgently needed to fully characterize the repertoire of antigen-specific TCRs.

To address these challenges, we developed a new approach based on the renewable nature of T cell libraries to screen for antigen-specific TCRs, exemplified in the context of patients with acute infectious mononucleosis (IM). Our method links natively paired TCRα:β gene sequences with their cognate pMHC targets. In addition, we measured the compound interaction affinities of individual TCRs, providing critical information on functional recognition needed to measure cross-reactivity in large pMHC panels. This new molecular platform will enable comprehensive functional studies of human T cell immunity in health and disease and accelerate the discovery of safe and effective immunotherapies based on TCRs.

## Results

### Single-cell cloning and sequencing of natively paired TCRα:β libraries

We previously described an emulsion-based technology for single-cell analysis of natively paired antibody heavy and light chains^30–33^, integrated with a cloning approach that allowed functional expression and screening of physically linked antibody heavy:light cDNAs^34–36^. Here, we modified these immune receptor display technologies to implement high-throughput screening and functional expression of natively paired TCRα:β sequences obtained from >10^6^ individual T cells per sample. T cells were obtained from peripheral blood mononuclear cells (PBMCs) of three acute infectious mononucleosis patients with acute IM (**Fig. 1, Supplementary Table 1**)^37,38^. A flow-focusing emulsification system was used to encapsulate single cells inside droplets containing poly-dT-coated magnetic beads to capture polyadenylated mRNAs. The magnetic beads were recovered, re-emulsified, and subjected to an overlap extension RT-PCR reaction physically linked TCRα and TCRβ amplicons onto the same cDNA strand (**Supplementary Fig. 1a, Supplementary Table 2**, see Methods)^30,32,39^. cDNA amplicons were analyzed by next-generation sequencing (NGS), and a nested PCR reaction was performed to generate TCRα:β amplicons for cloning into a new lentiviral expression vector, enabling library-scale display of the natively paired TCRα:β genes (**Fig. 1**, and **Supplementary Fig. 1b**).

**Figure 1.**
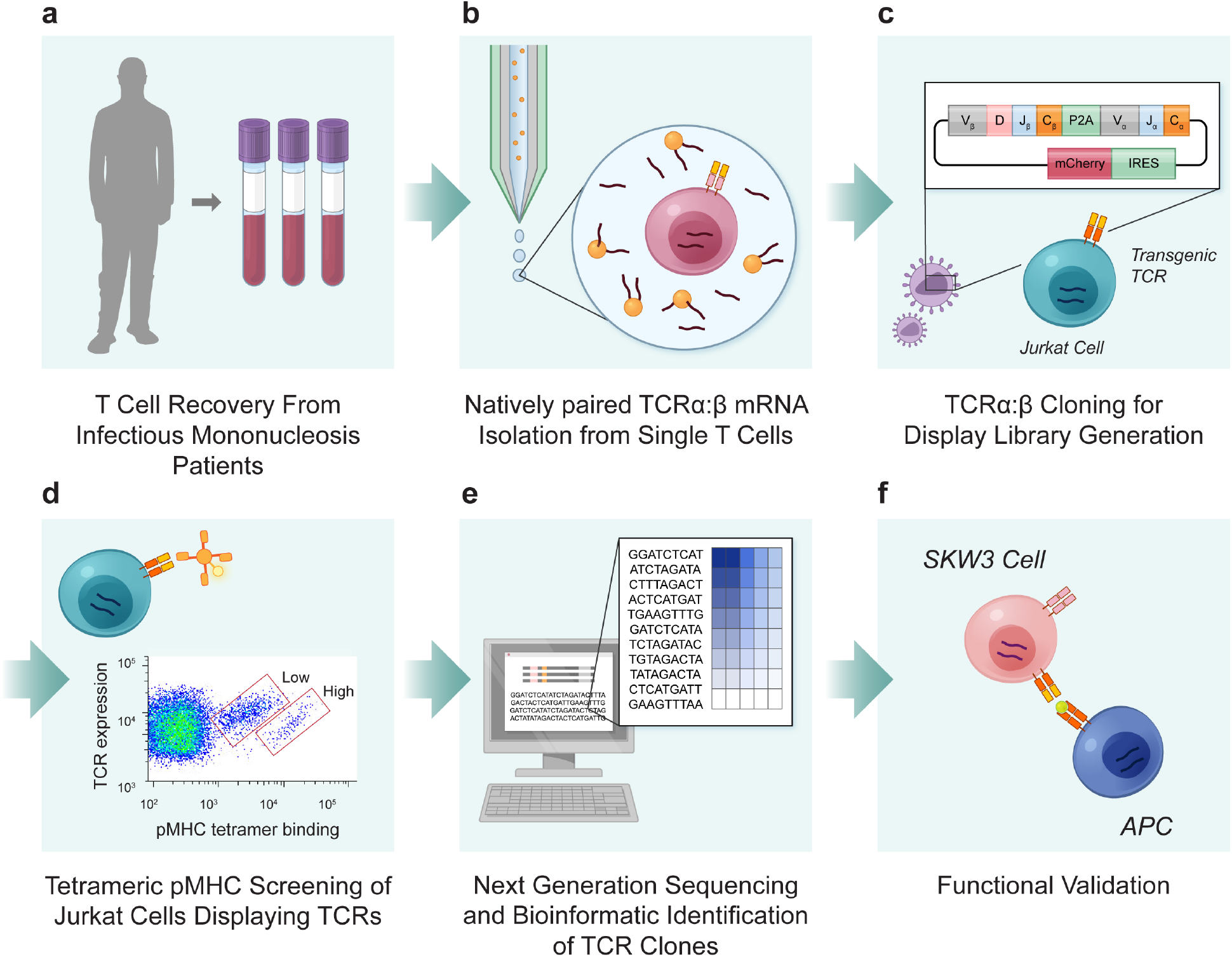
High-throughput TCRα:β sequencing and functional screening of human T cell repertoires. (**a**) PBMCs were collected from patients with acute IM, and T cells were expanded briefly *in vitro*. (**b**) Single T cells were isolated into emulsion droplets for mRNA capture, and an overlap extension RT-PCR was performed to physically link TCRβ variable region (*TRBV*) and TCRα variable region (*TRAV*) sequences onto the same DNA amplicon. (**c**) TCRα:β amplicons were cloned into a lentiviral display vector containing mCherry and IRES elements, followed by a 2A translation skip motif for complete TCRα:β expression. The TCRα:β plasmids were packaged into lentiviral particles to transduce Jurkat cells, creating immortalized TCRα:β libraries. (**d**) TCRα:β-Jurkat libraries were screened via FACS for binding to a panel of pMHC tetramers corresponding to epitopes derived from EBV. (**e**) TCR genes expressed by sorted Jurkat cells were characterized via NGS, and TCR sequences were tracked across sort rounds to determine the relative avidity and specificity of individual TCRs. (**f**) Monoclonal TCRs were transduced into SKW3 cells, and activation was assessed in response to pMHCs on the surface of antigen-presenting cells (APCs).

All three patients were seropositive for Epstein–Barr virus (EBV) at the time of sample acquisition (**Supplementary Table 1**). We began with an average of 10 × 10^6^ primary PBMCs from each patient, and following a brief period of *in vitro* stimulation, we analyzed TCRα:β amplicons via NGS and cloned the corresponding expression libraries into Jurkat cells (**Fig. 1a-c** and **Supplementary Tables 2–3**). Initial NGS analysis revealed 7,315,096 natively paired TCRα:β sequences. The data were then filtered for quality, clustered according to TCRβ chain nucleotide identity, and parsed to eliminate paired TCRα:β sequences with a read count of 1, leaving a total of 42,143 unique TCRα:β quality-filtered clusters across all three patients. Diverse TCR gene usage was observed in each library, confirming that large numbers of TCRs were recovered from each donor (**Fig. 2a, Supplementary Table 3**). As expected, small numbers of TCRβ sequences were shared among the repertories (**Fig. 2b**), each validated with a read count ≥2 to minimize the inclusion of amplification and sequencing errors in the final dataset^31–35^. Other recent studies included single-read TCRα:β sequences for repertoire-scale bioinformatic analyses^28^, and the corresponding outputs from our data are reported in the supplement to enable direct comparisons (**Supplementary Table 3**.)

**Figure 2.**
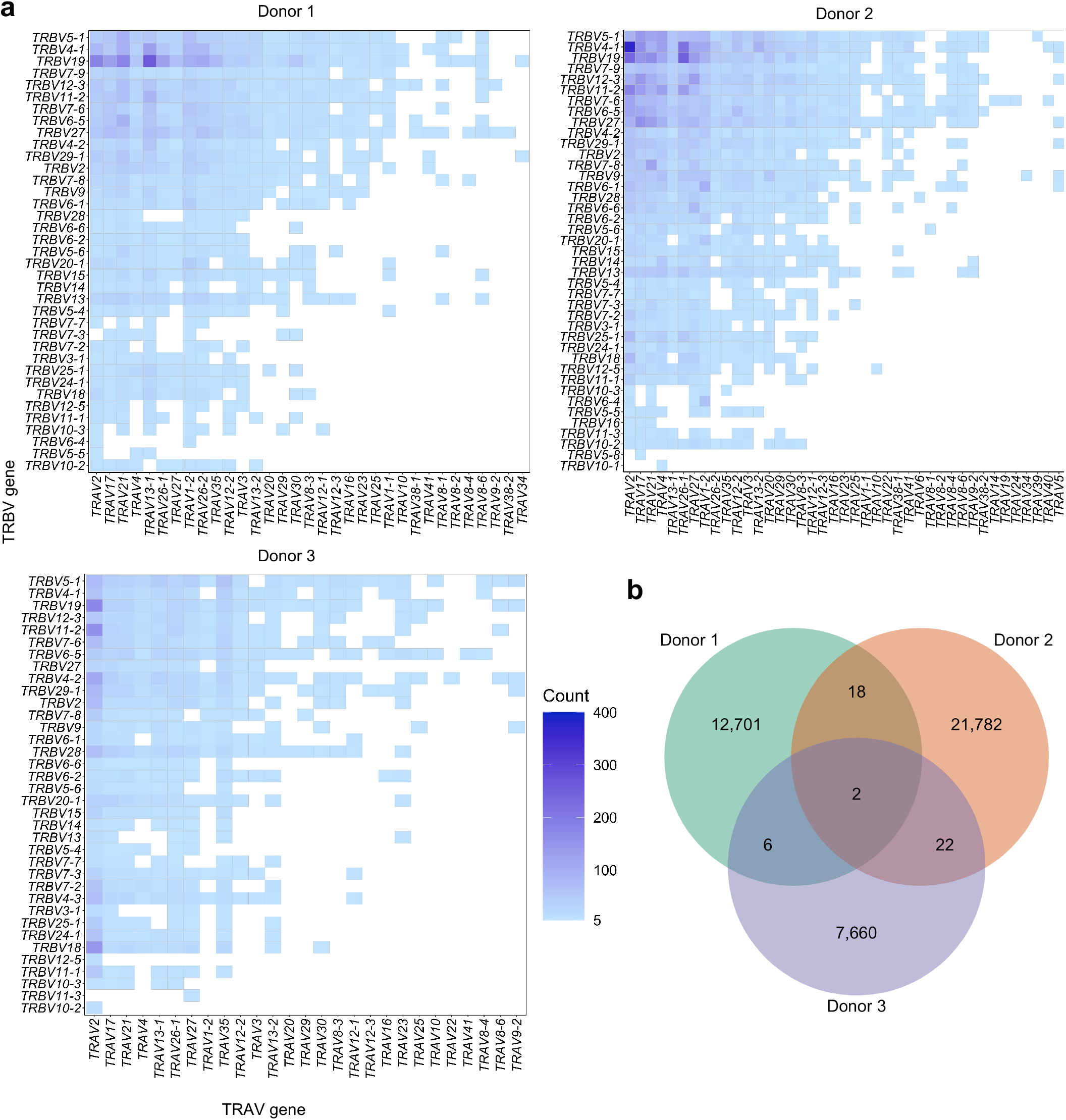
Genetic analysis of TCRα:β repertoires. (**a**) Paired *TRBV:TRAV* gene usage recovered from three patients with acute IM. (**b**) Genetic overlap of CDR-β3 nucleotide sequences among patients with IM. Shared clonotypes with a read difference >50-fold across samples were not included due to the potential for MiSeq index hopping during Illumina sequencing runs.

### Screening natively paired TCRα:β libraries against pMHC tetramers

We cloned TCRα:β amplicons into an immortalized surface display platform to enable library-scale interrogation of TCR binding to soluble pMHC antigens via fluorescence-activated cell sorting (FACS). Immortalized TCR libraries were screened using fluorescent pMHC tetramers, recovered via FACS, and analyzed by NGS to identify antigen-specific TCRs (**Fig 1d-e, Supplementary Fig. 1b**).

Paired TCRα:β amplicons were cloned into a lentiviral vector display system using an approach modified from previous studies^34–36^. The vector contained a leader sequence for TCRβ expression at one end of the restriction enzyme cut sites, with the required components of the TCRα constant region at the other end of the cut sites. We also included an internal ribosomal entry site (IRES) and an mCherry marker to identify TCRα:β expression via FACS (**Supplementary Fig. 1b**). After cloning the TCRα:β amplicons, the linker region was swapped with a linear construct containing the remaining portion of the TCRβ constant region, a sequence derived from porcine teschovirus-1 (P2A)^40^, and a leader sequence for TCRα expression (**Supplementary Fig. 1b**). At least 10^6^ transformants were maintained at each cloning step to preserve TCRα:β diversity^34–36^. The expression plasmids were packed into lentiviral particles for transduction into Jurkat cells, generating immortalized TCRα:β surface display libraries with the potential for limitless propagation.

We then leveraged the renewable nature of Jurkat cell libraries to screen the expressed TCRα:β repertories against multiple pMHCs. Each library was sorted for mCherry expression via FACS. mCherry^+^ TCRα:β-Jurkat libraries were then expanded and stained with a panel of EBV-derived pMHC antigens matched to the genetic background of each donor, using well-characterized fluorescent pMHC tetramers (**Supplementary Table 4**). In line with the diagnosis of acute IM, substantial reactivity was observed against EBV (**Fig. 3**). Cells labeled with fluorescent pMHC tetramers were collected, cultured, and sorted again to enrich the libraries for antigen-specific TCRs (**Fig. 3, Supplementary Fig. 2**). After two rounds of expansion, sorted cells were enriched 10–100-fold compared to the mCherry+ TCRα:β-Jurkat libraries (**Fig. 3, Supplementary Fig. 2**).

**Figure 3.**
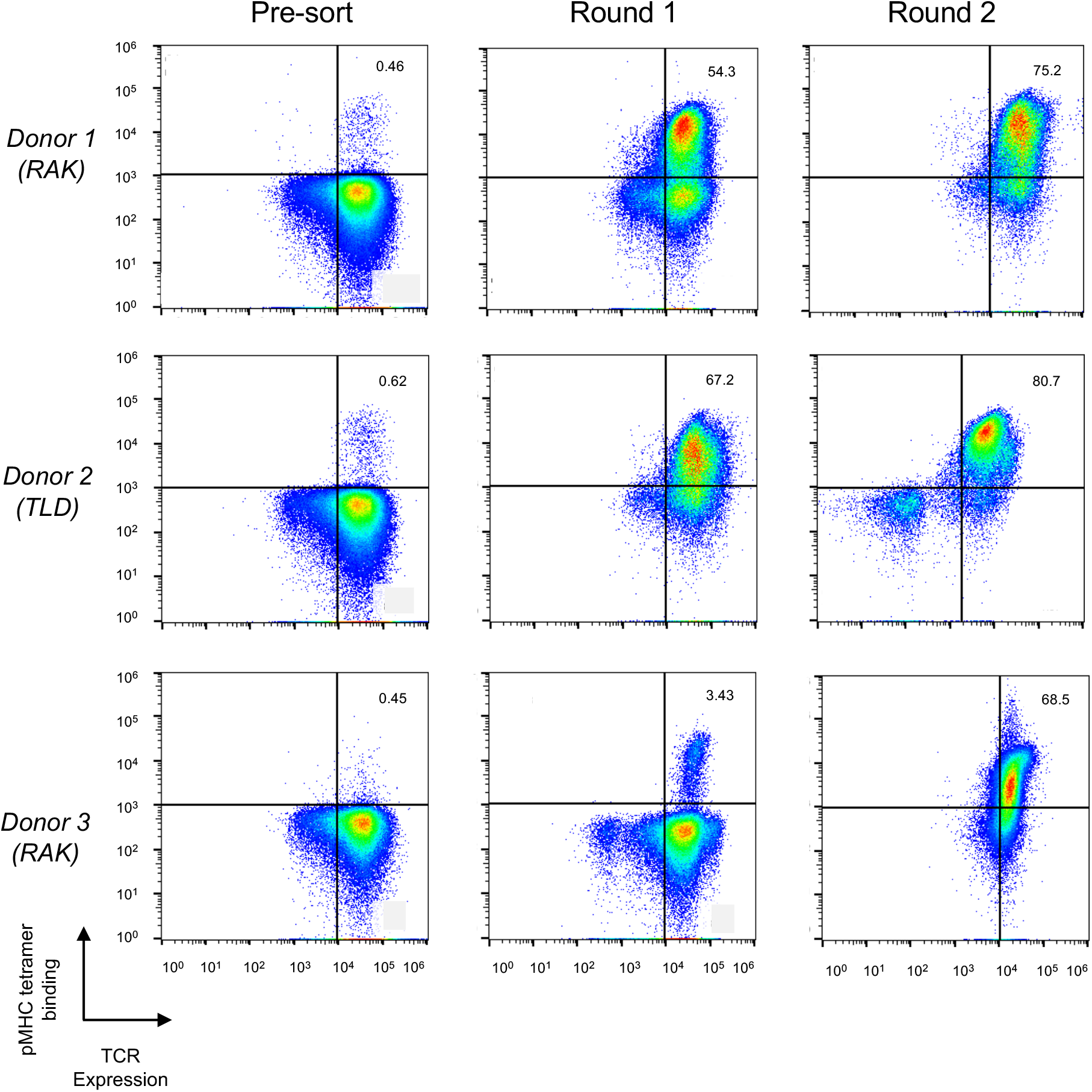
Functional screening of TCRα:β-Jurkat libraries against pMHC tetramers. (**a**–**c**) TCRα:β-Jurkat libraries were enriched via sequential rounds of FACS for TCRs that bound specific pMHCs. TCRα:β-Jurkat libraries were stained with pMHC tetramers corresponding to epitopes derived from EBV, namely TLD/HLA-A^*^02:01 and RAK/HLA-B^*^08:01. TCR expression is shown on the x-axis, and tetramer binding is shown on the y-axis.

### Identification of library-expressed TCRs specific for EBV

Expression libraries were sampled across each round of expansion and analyzed via NGS to quantify and track antigen-specific TCRs. We first picked single plasmid colonies after TCRα:β-Jurkat library amplification to validate our methodology. Single-colony analyses yielded seven unique clonotypes that bound to the TLD/HLA-A^*^02:01, QAK/HLA-B^*^08:01, or RAK/HLA-B^*^08:01 epitopes derived from EBV (**Table 1, Supplementary Table. 4**). The specificities of these clonotypes were maintained after expression as monoclonal TCRs (**Fig. 4a**).

**Table 1:**
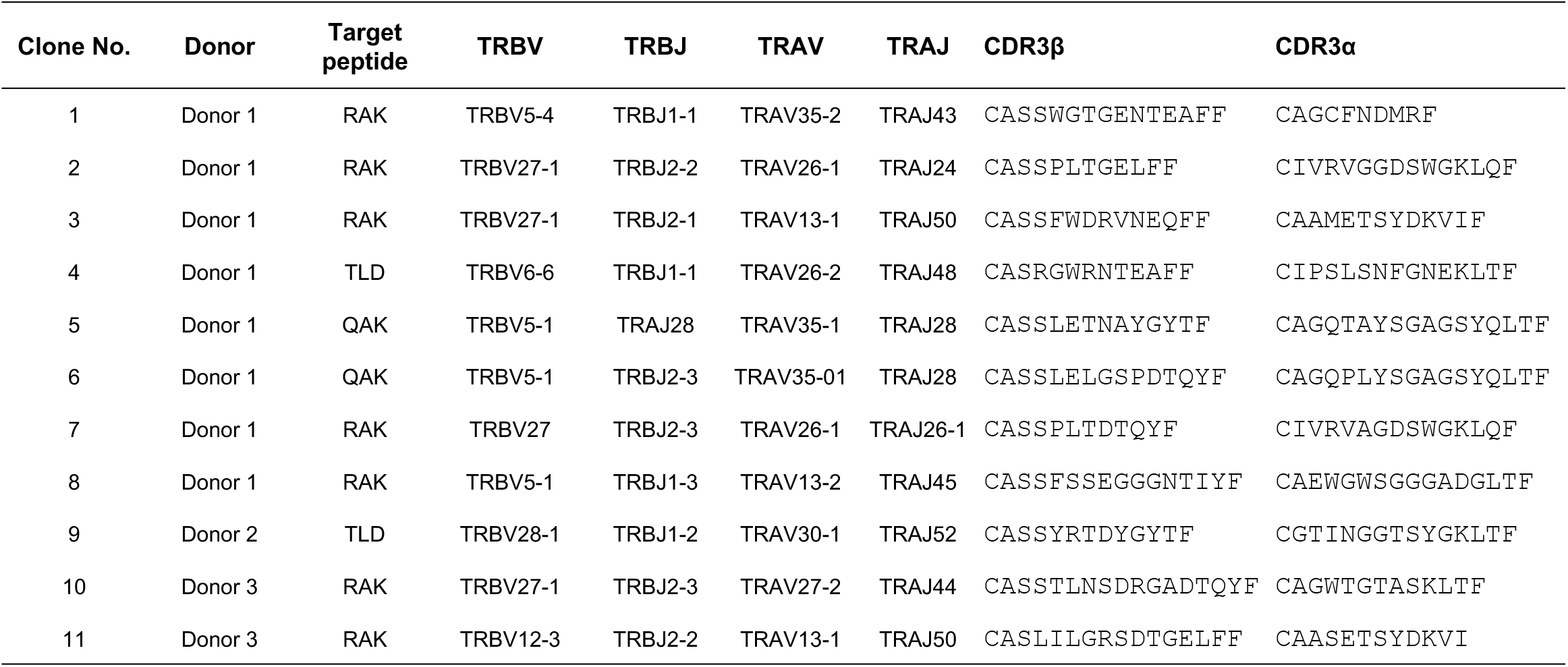
Genetic features of recovered TCRα:β clones

**Figure 4.**
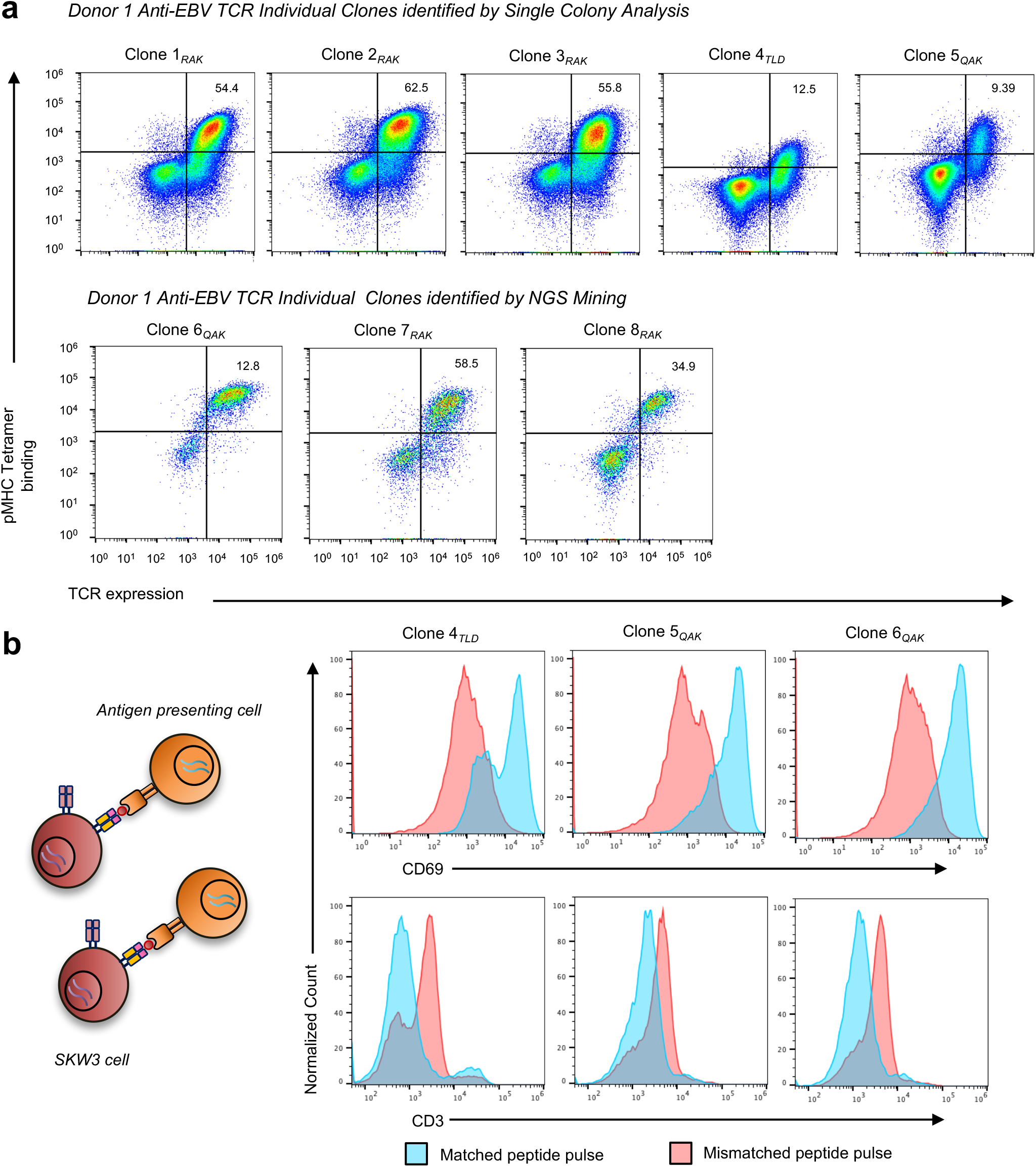
Functional validation of monoclonal TCRs recovered from high-throughput screening via NGS. (**a**) TCRα:β genes were transduced into Jurkat cells and validated for binding against cognate pMHCs. TCR expression is shown on the x-axis, and pMHC binding is shown on the y-axis. (**b**) Monoclonal TCRα:β-SKW3 cells were assessed for activation in response to pMHCs on the surface of antigen-presenting cells. CD69 or CD3 expression is shown on the x-axis, and normalized counts are shown on the y-axis.

We then analyzed the paired TCRα:β sequences in each library via NGS. Using this approach, we identified and validated an additional four EBV-specific TCRα:β clonotypes that had been enriched >30-fold after the final round of expansion. In particular, Clone 6 and Clone 8 bound to QAK/HLA-B^*^08:01, whereas Clone 7 and Clone 11 bound to RAK/HLA-B^*^08:01 (**Fig. 4a, Fig. 5b, Supplementary Table 4**). Of note, Clone 7 expressed a previously identified public TCRβ chain^3,41,42^, providing additional independent confirmation of TCR specificity.

**Figure 5.**
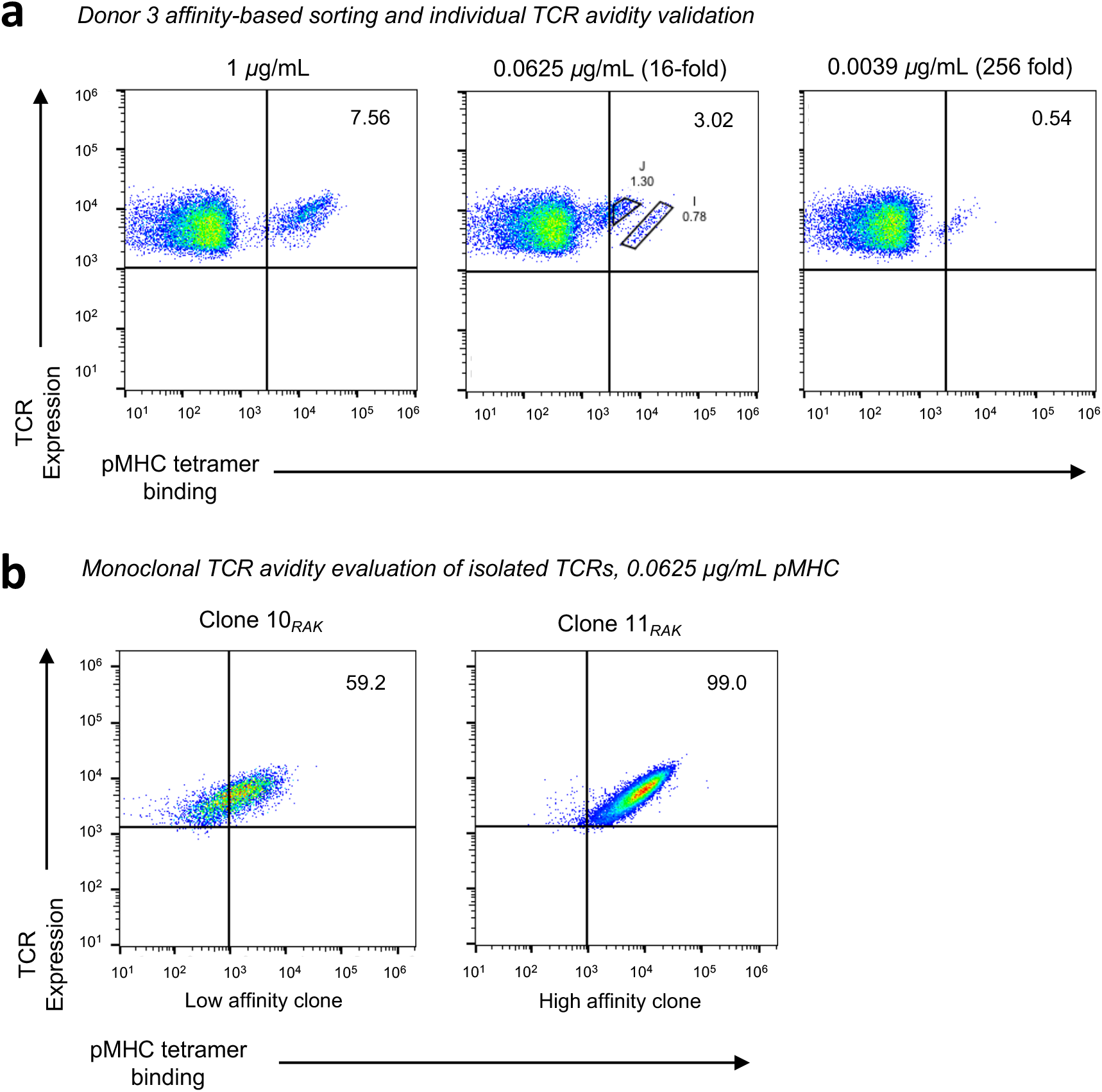
Avidity-based isolation of TCRα:β-Jurkat cells. (**a**) TCR avidity was assessed against of a gradient of pMHC tetramer concentrations, measuring TCR binding to the corresponding soluble antigens via FACS. Avidity-based sorts were performed from the Donor 3 library after 1 round of enrichment on the relevant pMHC. (**b**) Individual TCR clones identified in **a** were validated for pMHC binding at tetramer concentrations of 0.0625 µg/mL. Tetramer binding is shown on the x-axis, and TCR expression is shown on the y-axis.

### Functional validation of EBV-specific TCRs

To validate functional specificity of these clonotypes, we transduced the corresponding TCRα:β lentiviral particles into SKW3 cells. These reporter lines were then exposed to antigen-presenting cells expressing donor-matched HLA molecules pulsed with the relevant peptide. The transformed SKW3 cells were analyzed for surface expression of CD69, a common activation marker, and CD3, demarcating expression of the TCR^43,44^. As expected, TCR-SKW3 clones upregulated CD69 and downregulated CD3 after exposure to cognate pMHCs, and importantly, remained quiescent after exposure to non-cognate pMHCs (**Fig. 4b, Supplementary Fig. 3**). Overall, we found that 11/11 (100%) of the tested monoclonal TCRs that bound a cognate tetramer in the Jurkat cell display system also induced cellular activation in SKW3 cells exposed to the corresponding pMHC.

### Bulk techniques for assessing avidity of antigen-specific TCRs

TCR avidity is a translationally relevant parameter that identifies clonotypes with the capacity to recognize low densities of surface-expressed pMHCs. To evaluate such measurements in the context of our system, we stained the Donor 3 library with three different concentrations of the RAK/HLA-B^*^08:01 tetramer, ranging from 1 μg/mL down to 0.0039 μg/mL (**Fig. 5a**). Titration-based affinity assays fractionate the libraries based on differences in binding at various concentrations (**Fig. 5a**), and are commonly used to assess library-scale interactions^45,46^. Clear separation of the library into two distinct cell populations was observed at a tetramer concentration of 0.0625 μg/mL (**Fig. 5a**, *middle plot*), whereas only one cell population was observed at a tetramer concentration of 0.0039 μg/mL (**Fig. 5a**, *right plot*). Sequence analysis of these avidity-defined cell populations revealed the presence of two distinct monoclonal TCRs. Clone 10 was highly enriched in sort gate J (95%), indicating a low avidity for RAK/HLA-B^*^08:01, whereas Clone 11 was highly enriched in sort gate I (75.7%) and in the cell population identified at a tetramer concentration of 0.0039 μg/mL (77%), indicating a high avidity for RAK/HLA-B^*^08:01. The ability of our platform to support library-scale avidity measurements was further confirmed by expressing the sequences derived from Clone 10 and Clone 11 as monoclonal TCRs (**Fig. 5b, Supplementary Fig. 4**). These data confirmed that library-scale TCRα:β screening based on soluble pMHC titrations can compare the avidities of antigen-specific TCRs using bulk library-scale approaches paired with quantitative NGS data analysis.

## Discussion

Library-scale functional characterization of human TCRα:β specificity is a technically demanding task that is nonetheless critical for the advancement of clinical biotechnologies and for our understanding of human immunology. We developed and validated a new approach to this challenge based on TCR repertoire immortalization, iterative library screening against panels of pMHCs, and the functional characterization of monoclonal TCRs.

Our platform was adapted from previously reported strategies designed to link the functional and genetic features of natively paired antibody repertoires^34–36,47^, enabling a greater analytical throughput compared with previous single-cell studies of natively paired TCRs^4,20,48,49^. Of note, one microfluidic study demonstrated an ability to capture diverse clonotypes using a related approach and implemented two screening strategies to reduce the false-positive and false-negative discovery rates^28^. Here we identified an average of 14,047 unique TCRα:β sequences per sample, excluding singletons, which we believe are less reliable than clonotypes with 2 exact nucleotide-matched reads in datasets generated using NGS. Our data compared favorably with other high-throughput approaches incorporating functional analyses of TCRs^13,28^, and additional improvements in library efficiency and yield will be gained as the technology matures.

The ability to mine affinity-specificity relationships is clearly important for the rational design and discovery of protective TCRs^50,51^. Rapid affinity-based isolation procedures could also expedite the delivery of autologous immunotherapies with minimal off-target effects to enhnace drug efficacy and provide an inherent degree of safety. Our platform recovered individual antigen-specific TCR clonotypes with initial library frequencies as low as 0.00001, which were substantially enriched after purification via FACS. This remarkable ability to identify and reconstitute rare clonotypes with defined specificities has obvious advantages in the setting of personalized medicine. Additional modifications will further enhance our ability to interrogate functional TCRs. For example, the inclusion of oligonucleotide barcodes will enable massively parallel screening against hundreds of recombinant pMHCs^19^, and the generation of reporter libraries amenable to activation-based assays will allow the direct identification of functionally relevant TCRs, particularly for anti-viral and anti-cancer applications. In addition, the fidelity of library screening could be enhanced via CRISPR-based homology-directed recombination, limiting the expression profile of each transformed cell to a single TCR^10^. Our current approach has nonetheless addressed a major gap in functional TCRα:β profiling techniques, potentially enabling a greater understanding of adaptive immune responses and facilitating the development of more effective immunotherapeutic TCRs.

## Methods

### Study Participants

Donor 1 enrolled in a prospective study of primary EBV infection (University of Minnesota IRB 0608M90593). This participant developed IM at the age of 21, characterized by high fever, fatigue, body aches, and headache, with a maximum illness severity of 3, as reported previously^38^. Donors 2 and 3 enrolled in an experimental antiviral drug trial (University of Minnesota IRB 0709M16341). Donor 2 developed IM at the age of 21, characterized by fever, tender cervical lymph nodes, sore throat, and fatigue, with a maximum illness severity of 5. Donor 3 also developed IM at the age of 21, characterized by fever, loss of appetite, fatigue, sore throat, and headache, with a maximum illness severity of 3. Details of the trail were reported previously^52^. All donors provided written informed consent in accordance with the principles of the Declaration of Helsinki.

### Sample Collection and Handling

Donors provided oral wash samples by gargling with 22 mL of normal saline. Suspended oral cells were pelleted and frozen in two aliquots at -80 °C. Four aliquots of supernatant were also saved and frozen at -80 °C. PBMCs were isolated from venous blood samples via density gradient centrifugation over ACCUSPIN System-Histopaque-1077 (Sigma-Aldrich) and cryopreserved at 1 × 10^7^ cells/mL in heat-inactivated fetal bovine serum (Thermo Fisher Scientific) containing 10% dimethyl sulfoxide (Sigma-Aldrich). Cryopreserved PBMCs were thawed rapidly and diluted to 0.5 × 10^6^ cells/mL in complete CTS OpTmizer T Cell Expansion SFM (Thermo Fisher Scientific) containing 5% CTS Immune Cell SR (Thermo Fisher Scientific), 200 IU/mL IL-2 (National Cancer Institute Preclinical Biologics Repository), and 25 μL/mL ImmunoCult Human CD3/CD28 T Cell Activator (STEMCELL Technologies). Cells were expanded in RPMI 1640 medium (Thermo Fisher Scientific) containing 10% heat-inactivated fetal bovine serum (Thermo Fisher Scientific), 200 IU/mL IL-2 (National Cancer Institute Preclinical Biologics Repository), and 25 μL/mL ImmunoCult Human CD3/CD28 T Cell Activator (STEMCELL Technologies).

### Single Cell Emulsification

Expanded PBMCs were captured as single cells and emulsified for analyses of natively paired TCRα and TCRβ sequences using methods adapted from previous studies of antibody genes recovered from single B cells^30,32,34^. Briefly, a flow-focusing device was used to encapsulate individual T cells into emulsification droplets containing lysis buffer and oligo(dT) magnetic beads (New England Biolabs)^30^. Magnetic beads were suspended in lysis buffer, while cells were suspended in sterile phosphate buffered saline (PBS) at a concentration of 100,000 cells/mL. In addition to channels allowing for introduction of the bead/lysis buffer and cell/PBS mixtures, the flow-focusing device also contained a surrounding rapidly flowing oil phase comprising 4.5% (v/v) Span 80, 0.4% (v/v) Tween 80, 0.05% (v/v) Triton X-100, and 95.05% (v/v) light mineral oil (M5904, Sigma-Aldrich, St. Louis, MO). Single-cell emulsions generated by flow focusing were collected and stored on ice for 45 min. The aqueous droplets were then pooled and broken on ice using water-saturated diethyl ether (Fisher Scientific). Beads were re-emulsified in an overlap extension RT-PCR mixture using the SuperScript III RT-PCR Kit (Thermo Fisher Scientific, Waltham, MA) with the incorporation of custom designed TCR cloning primers (**Supplementary Table 2**)^53^. Natively paired TCRα and TCRβ sequences were physically linked via an overlap in the linker between the TRBC and TRAV regions. RT-PCR was performed under the following conditions: 30 min at 55 °C, 2 min at 94 °C, four cycles comprising 30 sec at 94 °C, 30 sec at 50 °C, and 2 min at 72 °C, four cycles comprising 30 sec at 94 °C, 30 sec at 55 °C, and 2 min at 72 °C, and 32 cycles comprising 30 sec at 94 °C, 30 sec at 60 °C, and 2 min at 72 °C, with a final hold for 7 min at 72 °C. RT-PCR products containing linked TCRα:β cDNA amplicons were purified using a DNA Clean & Concentrator Kit (D4033, Zymo Research, Irvine, CA).

### Semi-Nested PCR and Suppression PCR

A semi-nested PCR was performed using a HotStart GoTaq Polymerase System (M500,1 Promega, Madison, WI). In addition to the forward and reverse primers used to amplify the paired TCRα:β templates, a set of blocking oligonucleotides complementary to the 3’ region of the unfused TCRα and TCRβ products was designed with several nonsense nucleotides and a phosphate group at each 3’ end^54^. These nonsense regions were intended to suppress non-native pairing by causing a loss of homology between the elongated non-paired TCRα and TCRβ products. The first semi-nested PCR was performed under the following conditions: 2 min at 95 °C and 27 cycles comprising 35 sec at 95 °C, 40 sec at 58 °C, and 1 min at 73 °C, with a final hold for 5 min at 73 °C. The second semi-nested PCR was performed using a KAPA HiFi HotStart PCR Kit (7958897001, Roche, Basel, Switzerland) under the following conditions: 2 min at 95 °C and 15 cycles comprising 20 sec at 98 °C, 30 sec at 63 °C, and 30 sec at 72 °C, with a final hold for 7 min at 72 °C. PCR products were excised and purified from a 1.5% SYBR Safe Agarose Gel (Thermo Fisher Scientific).

### Natively Paired TCRα:β Library Cloning

The pLVX-EF1α-IRES-mCherry Vector (631987, Takara Bio, Mountain View, CA) was modified to enable surface expression of TCRs. After restriction enzyme digestion with BstBI and AgeI, the vector and the TCR amplicon were gel-purified and ligated using T4 DNA Ligase (M0202M, New England BioLabs Inc, Ipswich, MA) and the product was purified using a DNA Clean & Concentrator Kit (Zymo Research, Irvine, CA) and transformed via electroporation into competent MegaX DH10B T1 Electrocomp Cells (C640003, Invitrogen, Waltham, MA). Plasmids containing TCR libraries were purified using a ZymoPURE II Plasmid Maxiprep Kit (D4202, Zymo Research, Irvine, CA). The TRBC-p2A-TRAL insert was amplified using a KAPA HiFi HotStart PCR Kit (7958897001, Roche, Basel, Switzerland). Insert and the TCR library plasmids were digested with SpeI and Mlul, ligated as above with T4 DNA Ligase (New England BioLabs) and subsequent transformation and plasmid purification as reported above.

### HEK Cell Transfection and Lenti-X Concentration

HEK293 adherent cells grown to a confluency of 70–90% were transfected using a Lipofectamine 3000 Reagent Kit (L3000075, Invitrogen, Waltham, MA). Transfection procedures were adapted from the product manual. Briefly, 40 µL of Lipofectamine 3000 Reagent was diluted in 1 mL of Opti-MEM Reduced Serum Medium (31985062, Gibco, Waltham, MA) and a separate master mix was prepared comprising 1 mL Opti-MEM 8 µg PSPAX (12260, Addgene, Watertown, MA), 2 µg PMD2G plasmids (12259, Addgene), 40 µL of P3000 Reagent (Thermo Fisher Scientific), and 12 µg lentiviralpLVX-EF1α-IRES-mCherry plasmid for TCR expression per T75 flask (Thermo Fisher Scientific). This master mix was added to the diluted Lipofectamine 3000 Reagent and incubated for 15 min at room temperature. The final mixture was added dropwise to the flask containing HEK293 cells and incubated for 3 days at 37 °C. Viral supernatants from transfected HEK293 cells were collected and centrifuged at 800 × *g* for 10 min. For transduction of Jurkat and SKW3 cells, viral stocks were concentrated by adding supernatant at a 3:1 ratio to Lenti-X Concentrator (631232, Takara Bio, Mountain View, CA). Mixtures were incubated overnight at 4 °C. Samples were centrifuged at 1,500 × *g* for 45 min at 4 °C. Pellets were resuspended in 1 mL RPMI 1640 medium and stored at -80 °C. For transduction of primary T cell cells, viral supernatants were centrifuged at 25,000 rpm for 1.5 hr using an SW 28 Ti Rotor (Beckman Coulter, Brea, CA) re-suspended in 1.5 mL RPMI 1640 medium, and stored at -80 °C.

### TCR Transduction

For each transduction, 3 × 10^6^ Jurkat cells in 3 mL of RPMI 1640 medium were added to a single well of a 6-well plate, followed by 1 mL of the resuspended viral pellet and polybrene at a final concentration of 8 µg/mL (TR-1003, Sigma Aldrich, St. Louis, MO). After 24 hr, cells were pelleted via centrifugation at 500 × *g* for 8 min, resuspended in 10 mL of RPMI 1640 medium, transferred to T25 flasks, and incubated for 3 days at 37 °C. Cells were washed twice with PBS and sorted for mCherry expression via FACS.

### Tetramer Staining of Immortalized Cell Libraries

Fluorescent pMHC tetramers conjugated to BV421 were generated as described previously^55^, or purchased from The Tetramer Shop ApS (Kongens Lyngby, Denmark). Each stain incorporated 1 µg of tetramer, quantified with respect to the monomeric pMHC component, and 1 × 10^6^ TCRα:β-Jurkat cells in 100 µL of PBS containing 0.05% bovine serum albumin (BSA). Cells were incubated for 40 min at 37 °C and then washed three times in FACS buffer (PBS containing 0.05% BSA and 2 mM EDTA). Avidity-based screens were performed similarly via flow cytometry, using serial dilutions of each tetramer down to a concentration of 0.0039 μg/mL. In some experiments, cells were also stained with Alexa Fluor 488 anti-human TCR α/β (306712, BioLegend, San Diego, CA).

### Activation Assays in SKW3 Cells

Peptide pulse experiments were performed using several different antigen presenting cell lines (CIR-A2 and T2-B8), matched to donor HLA gene profiles. Individual TCR clones were cloned and packed as lentivirus as previously mentioned in Jurkat library generation, SKW3 cells were seeded at 0.6 × 10^5^ cells per monoclonal TCR in 3mL fresh RPMI media in 6-well plate. 400 μL of Lentivirus aliquot were add to SKW3 cells. Polybrene was added at 6μg/mL to enhance the transduction and incubated for 24 hrs. Cells were centrifuged at 500*g* × 5 mins, resuspended in 10 mL fresh RPMI media, and transferred into a T25 flask and incubated for 48 hrs. SKW3 cells were sorted at Day 3 and mCherry positive SKW3 cells were collected. Cells for each clone were expanded to 1 × 10^6^ cells/mL. APC cells (1 × 10^6^ cells) each of CIR-A2 and T2-B8 cells were transferred into 1.5 mL centrifuge tubes, washed with PBS twice, then resuspended with 100uL of FACS. APC CIR-A2 cells were pulsed with TLD peptide, while T2-B8 cells pulsed with either RAK or QAK peptides and incubated for 4 hrs at 4°C. APC cells were co-cultured with 1 mL of individual SKW3 clones (1 × 10^6^ cells) in 6-well plates and incubated overnight. Cells were harvested and cell activation were assessed by staining for CD69 (310904, BioLegend, San Diego, CA) and for CD3 (300426, BioLegend, San Diego, CA) and analyzed by flow cytometry.

### Bioinformatic Analysis

A previously described bioinformatic pipeline was adapted to identify natively paired TCRα:β sequences^30,32,34^. Modifications were incorporated to optimize the analysis of TCR genes. Raw sequences were quality-filtered and mapped to the *V, D*, and *J* genes, and CDR3 sequences were identified for each read using MiXCR v2.1.12 (ref. ^56^). Sequence data were filtered to exclude out-of-frame *V-(D)-J* combinations, and productive in-frame junction sequences were paired by Illumina read ID and compiled by CDR3 nucleotide and *V(D)J* gene identity. CDR-β3 nucleotide sequences were identified and clustered to 96% nucleotide sequence identity with terminal gaps ignored (USEARCH v5.2.32)^57^. We defined the set of TCRα:β clones recovered as all clusters with ≥2 reads in each data set. To determine the complete TCRα:β sequence, CDR3-α:β nucleotide sequences were used as anchors to map the germline TCRα and TCRβ genes with reference to the International ImMunoGeneTics (IMGT) Information System^58^.

## Acknowledgements

We thank David O. Schmeling for help with sample preparation and collection. This work was generously supported by grants from the Richard M. Schulze Family Foundation, the Randy Shaver Cancer Research and Community Fund, the Matt Cwiertny Foundation, the University of Minnesota Foundation, a Wellcome Trust Senior Investigator Award (100326/Z/12/Z, to D.A.P.), the University of Kansas Cancer Center, the US Department of Defense (W81XWH1810296), and the US National Institutes of Health (DP5OD023118, P20GM103638, P20GM103418, and R21CA230487).

## Ethics statement

Clinical trials were approved by the University of Minnesota Institutional Review Board. Informed consent with permission to use stored samples for subsequent research investigations was obtained from all donors in accordance with the principles of the Declaration of Helsinki.

## Data Availability

Raw sequence data were deposited in the NCBI Sequence Read Archive (SRA) under accession number XXXXXXX.

---

**Supplementary Figure 1.**
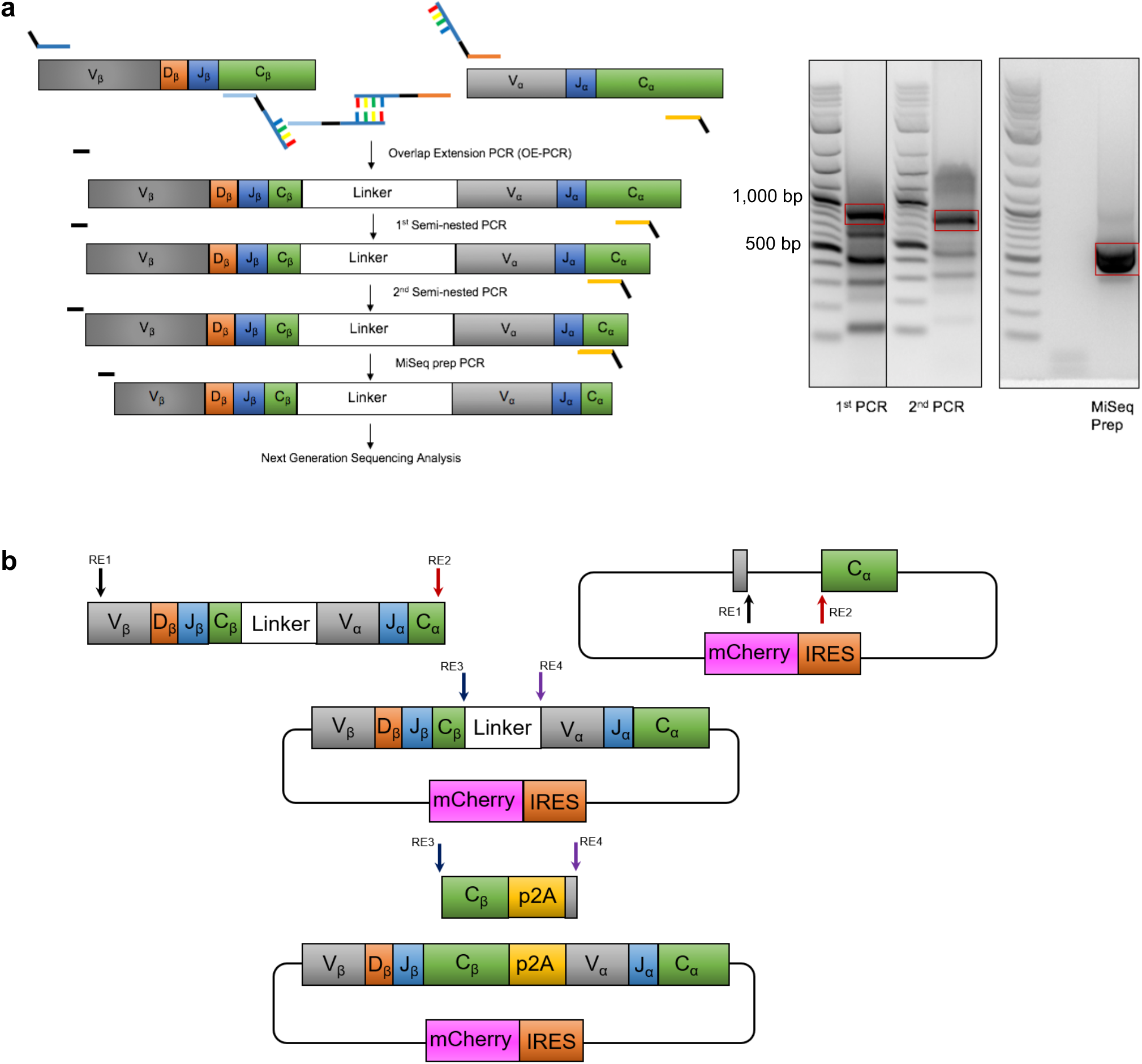
(**a**) Overlap extension RT-PCR of natively paired TCRα:β amplicons. (**b**) Cloning strategy for amplified TCRs. The overlap extension RT-PCR linker sequence was substituted with a p2A expression cassette for lentiviral transduction of TCRα:β libraries.

**Supplementary Figure 2.**
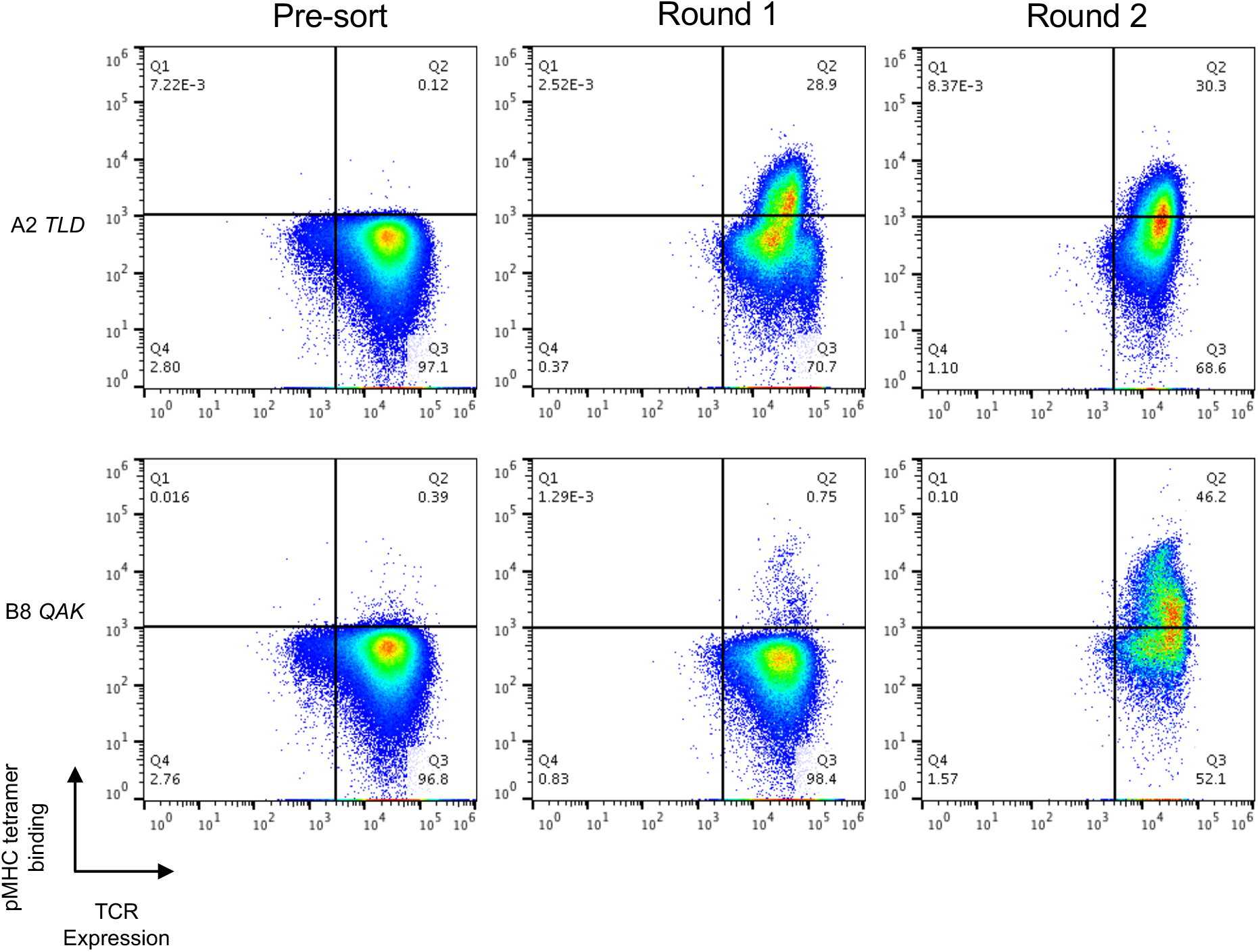
The TCRα:β-Jurkat library from Donor 1 was screened for pMHC tetramer binding via sequential rounds of FACS. U*pper panel*, TLD/HLA-A^*^02:01; *lower panel*, QAK/HLA-B^*^08:01. TCR expression is shown on the x-axis, and tetramer binding is shown on the y-axis.

**Supplementary Figure 3.**
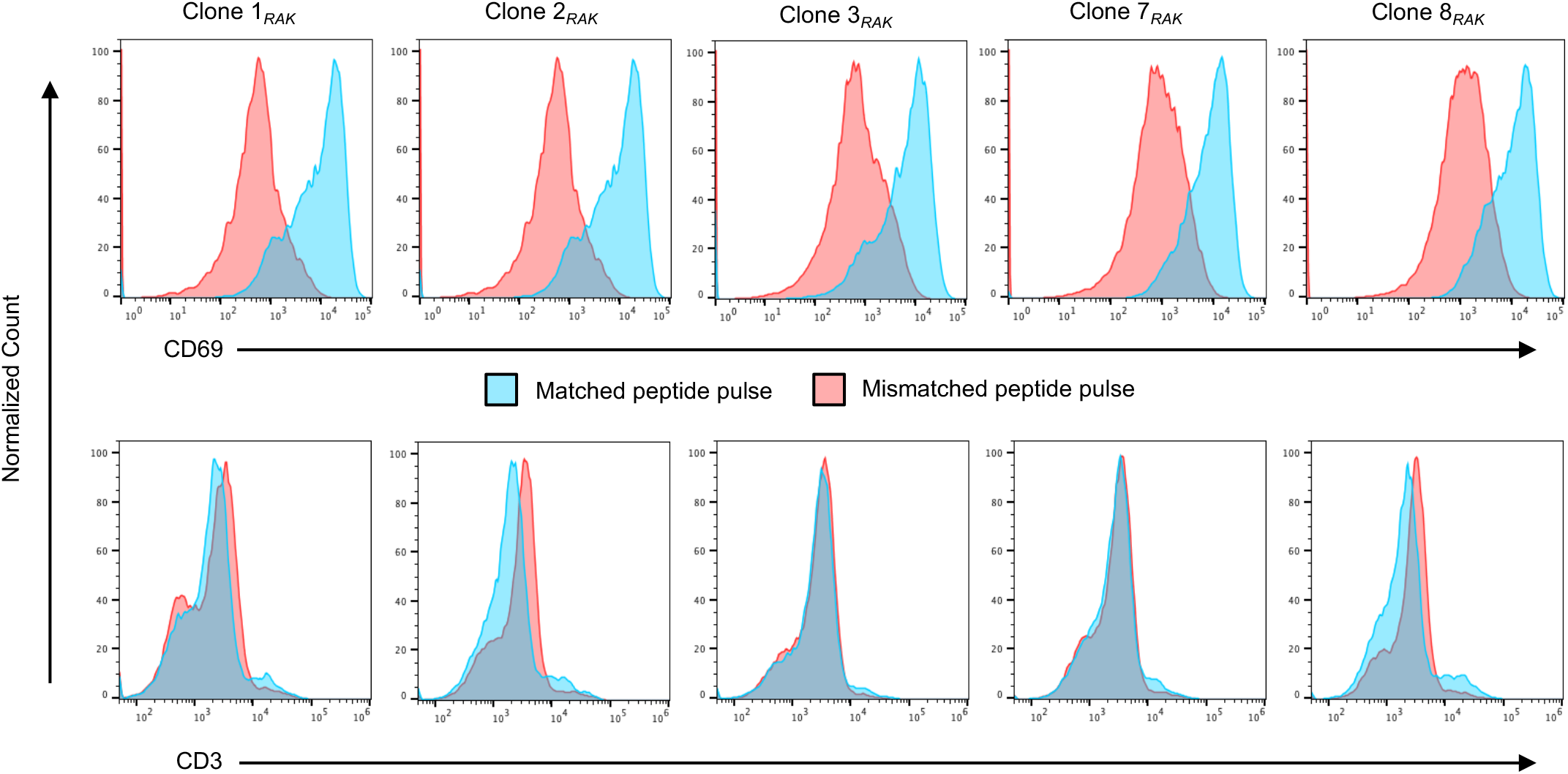
Flow cytometric analyses of monoclonal TCRs expressed in SKW3 cells after exposure to pMHCs on the surface of antigen-presenting cells. CD69 or CD3 expression is shown on the x-axis, and normalized counts are shown on the y-axis.

**Supplementary Figure 4.**
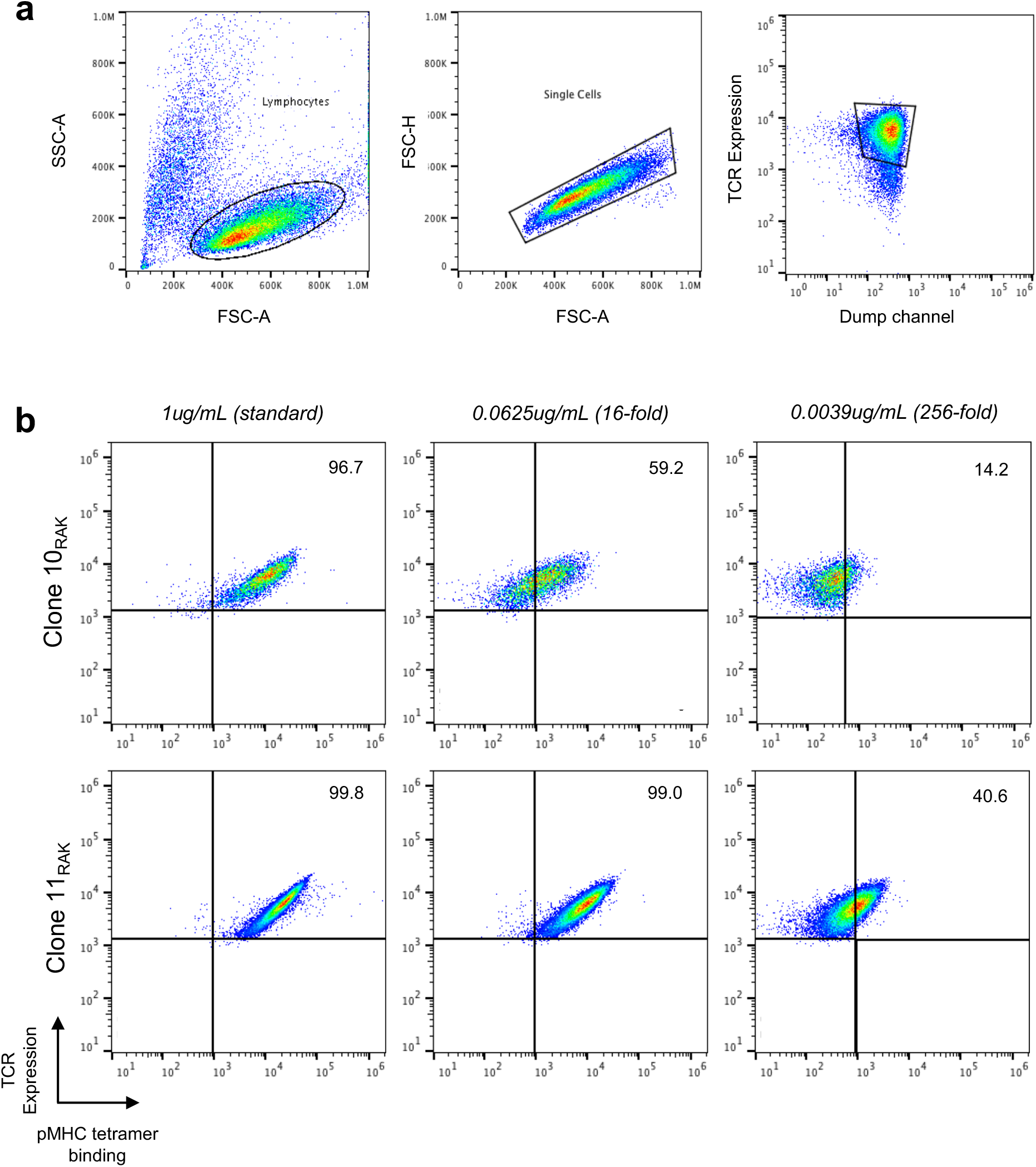
(**a**) Gating strategy for avidity-based assessments via FACS. Cells were selected on the basis of CD3 expression before experimentation. (**b**) Avidity-based assessments for monoclonal TCRα:β Clone 10_RAK_ (upper panel) and monoclonal TCRα:β Clone 11_RAK_ (lower panel) were performed using the RAK/HLA-B^*^08:01 tetramer at concentrations of 1 µg/mL, 0.0625 µg/mL, and 0.0039 µg/mL.

**Supplementary Table 1.** Donor information for EBV samples used in this study.

| Patient ID | HLA*A2 | HLA*B8 |
| --- | --- | --- |
| Donor 1 | Yes | Yes |
| Donor 2 | Yes | No |
| Donor 3 | Yes | Yes |

**Supplementary Table 2.**
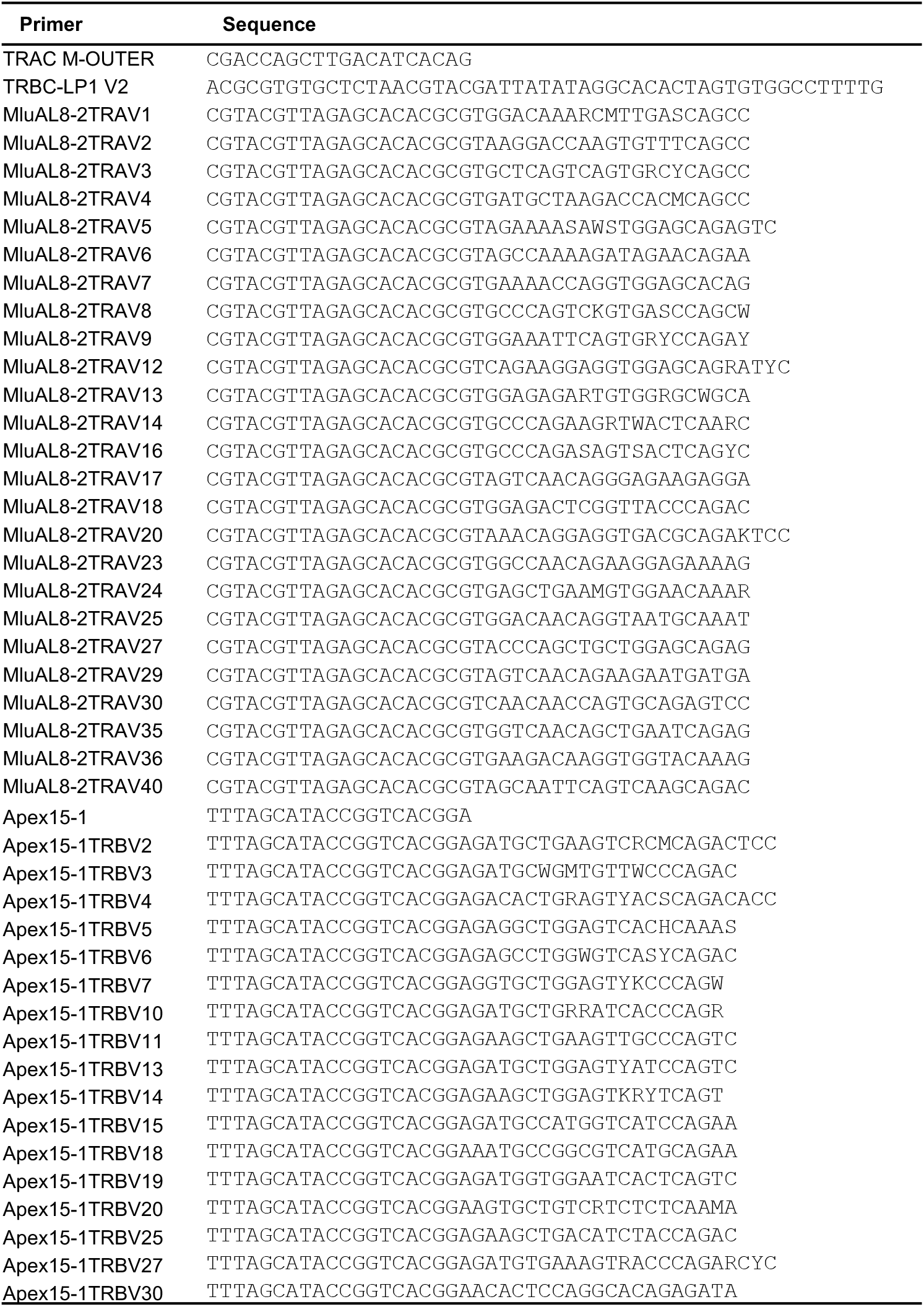
Primers used for OE RT-PCR amplification

**Supplementary Table 3.** Key statistics for the next generation sequencing analysis of patient T cell receptor repertoires.

| Sample Name | No. Expanded<br>PBMCs Analyzed | Number of Raw<br>Reads | Reads Passing<br>Quality Filter | Quality-filtered<br>TCRα:β<br>Sequences | Observed<br>TCRα:β Clusters<br>with ≥1 Reads | Observed<br>TCRα:β Clusters<br>with ≥2 Reads |
| --- | --- | --- | --- | --- | --- | --- |
| Donor 1 | 3.5×10<sup>6</sup> | 3,165,367 | 3,161,379 | 2,894,718 | 267,127 | 12,701 |
| Donor 2 | 1.5×10<sup>6</sup> | 7,081,113 | 7,070,723 | 2,927,183 | 312,740 | 21,782 |
| Donor 3 | 4.0×10<sup>6</sup> | 3,214,081 | 3,210,746 | 1,493,195 | 147,907 | 7,660 |

**Supplementary Table 4.** EBV tetramer sorting probes used in the study.

| EBV Gene | MHC Allele | Peptide Sequence |
| --- | --- | --- |
| *BMLF1* | HLA-A*02:01 | GLCTLVAML |
| *BMRF1* | HLA-A*02:01 | TLDYKPLSV |
| *BRLF1* | HLA-A*02:01 | YVLDHLIVV |
| *EBNA3A* | HLA-B*08:01 | QAKWRLQTL |
| *BZLF1* | HLA-B*08:01 | RAKFKQLL |
| *EBNA3A* | HLA-B*08:01 | FLRGRAYGL |

## References

1. Krogsgaard, M. & Davis, M. M. How T cells ‘see’ antigen. Nat Immunol 6, 239–245 (2005).

2. Sewell, A. K. Why must T cells be cross-reactive? Nat Rev Immunol 12, 669–677 (2012).

3. Koning, D. et al. In vitro expansion of antigen-specific CD8+ T cells distorts the T-cell repertoire. Journal of Immunological Methods 405, 199–203 (2014).

4. Linnemann, C. et al. High-throughput identification of antigen-specific TCRs by TCR gene capture. Nat Med 19, 1534–1541 (2013).

5. Scheper, W. et al. Low and variable tumor reactivity of the intratumoral TCR repertoire in human cancers. Nat Med 25, 89–94 (2019).

6. Yossef, R. et al. Enhanced detection of neoantigen-reactive T cells targeting unique and shared oncogenes for personalized cancer immunotherapy. JCI Insight 3, e122467 (2018).

7. Hu, Z. et al. A cloning and expression system to probe T-cell receptor specificity and assess functional avidity to neoantigens. Blood 132, 1911–1921 (2018).

8. King, C. G. et al. T Cell Affinity Regulates Asymmetric Division, Effector Cell Differentiation, and Tissue Pathology. Immunity 37, 709–720 (2012).

9. Aleksic, M. et al. Dependence of T Cell Antigen Recognition on T Cell Receptor-Peptide MHC Confinement Time. Immunity 32, 163–174 (2010).

10. Vazquez-Lombardi, R. et al. CRISPR-targeted display of functional T cell receptors enables engineering of enhanced specificity and prediction of cross-reactivity. http://biorxiv.org/lookup/doi/10.1101/2020.06.23.166363 (2020) doi:10.1101/2020.06.23.166363.

11. Szabo, P. A. et al. Single-cell transcriptomics of human T cells reveals tissue and activation signatures in health and disease. Nat Commun 10, 4706 (2019).

12. Corridoni, D. et al. Single-cell atlas of colonic CD8+ T cells in ulcerative colitis. Nat Med 26, 1480–1490 (2020).

13. Zhang, Z., Xiong, D., Wang, X., Liu, H. & Wang, T. Mapping the functional landscape of T cell receptor repertoires by single-T cell transcriptomics. Nat Methods 18, 92–99 (2021).

14. Newell, E. W. & Davis, M. M. Beyond model antigens: high-dimensional methods for the analysis of antigen-specific T cells. Nat Biotechnol 32, 149–157 (2014).

15. Ornatsky, O., Baranov, V. I., Bandura, D. R., Tanner, S. D. & Dick, J. Multiple cellular antigen detection by ICP-MS. Journal of Immunological Methods 308, 68–76 (2006).

16. Stronen, E. et al. Targeting of cancer neoantigens with donor-derived T cell receptor repertoires. Science 352, 1337–1341 (2016).

17. Glanville, J. et al. Identifying specificity groups in the T cell receptor repertoire. Nature 547, 94–98 (2017).

18. Bentzen, A. K. et al. Large-scale detection of antigen-specific T cells using peptide-MHC-I multimers labeled with DNA barcodes. Nat Biotechnol 34, 1037–1045 (2016).

19. Zhang, S.-Q. et al. High-throughput determination of the antigen specificities of T cell receptors in single cells. Nat Biotechnol 36, 1156–1159 (2018).

20. Nesterenko, P. A. et al. Droplet-based mRNA sequencing of fixed and permeabilized cells by CLInt-seq allows for antigen-specific TCR cloning. Proc Natl Acad Sci USA 118, e2021190118 (2021).

21. Zhang, J.-Y. et al. Single-cell landscape of immunological responses in patients with COVID-19. Nat Immunol 21, 1107–1118 (2020).

22. Azizi, E. et al. Single-Cell Map of Diverse Immune Phenotypes in the Breast Tumor Microenvironment. Cell 174, 1293-1308.e36 (2018).

23. Gantner, P. et al. Single-cell TCR sequencing reveals phenotypically diverse clonally expanded cells harboring inducible HIV proviruses during ART. Nat Commun 11, 4089 (2020).

24. Mateus, J. et al. Selective and cross-reactive SARS-CoV-2 T cell epitopes in unexposed humans. Science 370, 89–94 (2020).

25. Grant, E. J. et al. Broad CD8+ T cell cross-recognition of distinct influenza A strains in humans. Nat Commun 9, 5427 (2018).

26. Mendoza, J. L. et al. Interrogating the recognition landscape of a conserved HIV-specific TCR reveals distinct bacterial peptide cross-reactivity. eLife 9, e58128 (2020).

27. Linette, G. P. et al. Cardiovascular toxicity and titin cross-reactivity of affinity-enhanced T cells in myeloma and melanoma. Blood 122, 863–871 (2013).

28. Spindler, M. J. et al. Massively parallel interrogation and mining of natively paired human TCRαβ repertoires. Nat Biotechnol 38, 609–619 (2020).

29. Guo, X. J. et al. Rapid cloning, expression, and functional characterization of paired αβ and γδ T-cell receptor chains from single-cell analysis. Molecular Therapy - Methods & Clinical Development 3, 15054 (2016).

30. DeKosky, B. J. et al. In-depth determination and analysis of the human paired heavy- and light-chain antibody repertoire. Nat Med 21, 86–91 (2015).

31. DeKosky, B. J. et al. Large-scale sequence and structural comparisons of human naive and antigen-experienced antibody repertoires. Proc Natl Acad Sci USA 113, E2636–E2645 (2016).

32. McDaniel, J. R., DeKosky, B. J., Tanno, H., Ellington, A. D. & Georgiou, G. Ultra-high-throughput sequencing of the immune receptor repertoire from millions of lymphocytes. Nat Protoc 11, 429–442 (2016).

33. Lagerman, C. E. et al. Ultrasonically-guided flow focusing generates precise emulsion droplets for high-throughput single cell analyses. Journal of Bioscience and Bioengineering 128, 226–233 (2019).

34. Wang, B. et al. Functional interrogation and mining of natively paired human VH:VL antibody repertoires. Nat Biotechnol 36, 152–155 (2018).

35. Fahad, A. S. et al. Functional Profiling of Antibody Immune Repertoires in Convalescent Zika Virus Disease Patients. Front. Immunol. 12, 615102 (2021).

36. Banach, B. B. et al. Paired heavy and light chain signatures contribute to potent SARS-CoV-2 neutralization in public antibody responses. http://biorxiv.org/lookup/doi/10.1101/2020.12.31.424987 (2021) doi:10.1101/2020.12.31.424987.

37. Grimm, J. M. et al. Prospective studies of infectious mononucleosis in university students. Clin Trans Immunol 5, e94 (2016).

38. Balfour, H. H. et al. Behavioral, Virologic, and Immunologic Factors Associated With Acquisition and Severity of Primary Epstein–Barr Virus Infection in University Students. The Journal of Infectious Diseases 207, 80–88 (2013).

39. DeKosky, B. J. et al. High-throughput sequencing of the paired human immunoglobulin heavy and light chain repertoire. Nat Biotechnol 31, 166–169 (2013).

40. Liu, Z. et al. Systematic comparison of 2A peptides for cloning multi-genes in a polycistronic vector. Sci Rep 7, 2193 (2017).

41. Miconnet, I. et al. Large TCR Diversity of Virus-Specific CD8 T Cells Provides the Mechanistic Basis for Massive TCR Renewal after Antigen Exposure. J.I. 186, 7039–7049 (2011).

42. Koning, D. et al. CD8 ^+^ TCR Repertoire Formation Is Guided Primarily by the Peptide Component of the Antigenic Complex. J.I. 190, 931–939 (2013).

43. Nguyen, T. H. O. et al. Recognition of Distinct Cross-Reactive Virus-Specific CD8 ^+^ T Cells Reveals a Unique TCR Signature in a Clinical Setting. J.I. 192, 5039–5049 (2014).

44. Mayassi, T. et al. Chronic Inflammation Permanently Reshapes Tissue-Resident Immunity in Celiac Disease. Cell 176, 967-981.e19 (2019).

45. Starr, T. N. et al. Deep Mutational Scanning of SARS-CoV-2 Receptor Binding Domain Reveals Constraints on Folding and ACE2 Binding. Cell 182, 1295-1310.e20 (2020).

46. Harris, D. T. et al. Deep Mutational Scans as a Guide to Engineering High Affinity T Cell Receptor Interactions with Peptide-bound Major Histocompatibility Complex. Journal of Biological Chemistry 291, 24566–24578 (2016).

47. Madan, B. et al. Mutational fitness landscapes reveal genetic and structural improvement pathways for a vaccine-elicited HIV-1 broadly neutralizing antibody. Proc Natl Acad Sci USA 118, e2011653118 (2021).

48. Trautmann, L. et al. Selection of T Cell Clones Expressing High-Affinity Public TCRs within Human Cytomegalovirus-Specific CD8 T Cell Responses. J Immunol 175, 6123–6132 (2005).

49. Day, E. K. et al. Rapid CD8 ^+^ T Cell Repertoire Focusing and Selection of High-Affinity Clones into Memory Following Primary Infection with a Persistent Human Virus: Human Cytomegalovirus. J Immunol 179, 3203–3213 (2007).

50. Campillo-Davo, D., Flumens, D. & Lion, E. The Quest for the Best: How TCR Affinity, Avidity, and Functional Avidity Affect TCR-Engineered T-Cell Antitumor Responses. Cells 9, 1720 (2020).

51. Schmid, D. A. et al. Evidence for a TCR Affinity Threshold Delimiting Maximal CD8 T Cell Function. J.I. 184, 4936–4946 (2010).

52. Balfour, H. et al. Randomized, Placebo-controlled, Double-blind Trial of Valomaciclovir (VALM) for Infectious Mononucleosis (IM). in V-1256a (Prohealth, 2009).

53. Boria, I., Cotella, D., Dianzani, I., Santoro, C. & Sblattero, D. Primer sets for cloning the human repertoire of T cell Receptor Variable regions. BMC Immunol 9, 50 (2008).

54. Turchaninova, M. A. et al. Pairing of T-cell receptor chains via emulsion PCR: New technology. Eur. J. Immunol. 43, 2507–2515 (2013).

55. Price, D. A. et al. Avidity for antigen shapes clonal dominance in CD8+ T cell populations specific for persistent DNA viruses. Journal of Experimental Medicine 202, 1349–1361 (2005).

56. Bolotin, D. A. et al. MiXCR: software for comprehensive adaptive immunity profiling. Nature Methods 12, 380–381 (2015).

57. Edgar, R. C. Search and clustering orders of magnitude faster than BLAST. Bioinformatics 26, 2460–2461 (2010).

58. Lefranc, M.-P. et al. IMGT®, the international ImMunoGeneTics information system® 25 years on. Nucleic Acids Research 43, D413–D422 (2015).

